# m6A-ELISA, a simple method for quantifying *N6*-methyladenosine from mRNA populations

**DOI:** 10.1101/2022.09.27.509679

**Authors:** Imke Ensinck, Theodora Sideri, Miha Modic, Charlotte Capitanchik, Patrick Toolan-Kerr, Folkert J. van Werven

**Affiliations:** The Francis Crick Institute, 1 Midland Road, London, NW1 1AT, UK

## Abstract

*N6*-methyladenosine (m6A) is a widely studied and abundant RNA modification. The m6A mark regulates the fate of RNAs in various ways, which in turn, drives changes in cell physiology, development, and disease pathology. Over the last decade, numerous methods have been developed to map and quantify m6A sites genomewide through deep sequencing. Alternatively, m6A levels can be quantified from a population of RNAs using techniques such as liquid chromatography-mass spectrometry or thin layer chromatography. However, many methods for quantifying m6A levels involve extensive protocols and specialized data analysis, and often only a few samples can be handled in a single experiment. Here, we developed a simple method for determining m6A levels in mRNA populations from various sources based on enzyme-linked immunosorbent-based assay (m6A-ELISA). We have optimized various steps of m6A-ELISA such as sample preparation and the background signal resulting from the primary antibody. We validated the method using mRNA populations from budding yeast and mouse embryonic stem cells. The full protocol takes less than a day, requiring only 25 ng of mRNA. The m6A-ELISA protocol is therefore quick, cost-effective, and scalable, making it a valuable tool for determining relative m6A levels in samples from various sources that could be adapted to detect other mRNA modifications.

## Introduction

Epitranscriptomics, the study of posttranscriptional base modifications of RNAs, has been an emerging field of study for the last decade. Among all RNA modifications, *N6*-methyladenosine (m6A) is one of the most widespread and widely studied. Writer and reader proteins of the m6A RNA modification exert numerous functions in controlling the fate of mRNAs in eukaryotes, and play critical roles in development, differentiation and disease pathology (Zaccara et al. 2019; Yang et al. 2020; He and He 2021). Importantly, the levels of m6A are highly dynamic between species, cell types, and conditions (Schwartz et al. 2013a; Roignant and Soller 2017; Yang et al. 2018). Hence, techniques for measuring m6A levels are essential for providing insights on the abundance and dynamics of m6A containing RNAs.

Various methods have been developed to measure m6A from mRNA populations, within single transcripts, and at nucleotide resolution (Linder et al. 2015; Bodi and Fray 2017; Garcia-Campos et al. 2019; Dierks et al. 2021; Leger et al. 2021; Mirza et al. 2022). Each of these has their purpose in helping to understand the various aspects of m6A biology. Currently only a few techniques are available for determining m6A levels in RNA populations. These include thin layer chromatography (TLC), m6A RNA dot blot, and mass-spectrometry (MS) of RNA fragmented into nucleosides (Bodi and Fray 2017; Nagarajan et al. 2019; Mathur et al. 2021). Techniques such as TLC and m6A RNA dot blot are relatively time consuming and low throughput. Additionally, measuring RNA modifications by MS often requires access to specialized instruments and specialized training. Therefore, simple and rapid techniques for measuring m6A levels in RNA samples would be useful for m6A researchers and the epitranscriptomics field as a whole.

Here we present an indirect enzyme-linked immunosorbent assay for the detection of m6A (m6A-ELISA), a method for measuring relative changes in m6A levels across mRNA samples. We optimized several steps in the protocol to obtain a high signal-to-noise ratio using yeast mRNAs. Furthermore, we show that the method can detect dynamic changes in m6A levels in yeast and relative m6A levels in mouse embryonic stem cells (ESCs). The m6A-ELISA protocol is simple, cost efficient, and can potentially be adopted for studying other RNA modifications.

## Results and discussion

### m6A levels can be detected in mRNA using an ELISA-like method

To measure m6A levels within an RNA population, we set out to develop a detection method based on ELISA (Lin 2015). In short, mRNA is bound directly to a microplate using a commercially available nucleic acid microplate binding solution. The bound mRNA is then incubated with a primary anti-m6A antibody. This incubation is followed by the addition of a secondary, HRP-coupled antibody which allows a colorimetric readout using generic ELISA substrates.

To optimize the m6A-ELISA, we considered variables that could influence the signal-to-noise ratio. These variables included background signal from non-specific binding by primary antibodies, blocking reagents, and the method of RNA preparation. We also generated control samples to allow the use of a standard curve. Tests were performed with mRNA isolated from diploid budding yeast cells in the early phase of the meiotic program. In this stage, m6A is abundant because the m6A writer complex, including the yeast METLL3 orthologue Ime4, is expressed and active (Clancy et al. 2002; Agarwala et al. 2012). Importantly, diploid cells harbouring an *IME4* gene deletion display no detectable levels of m6A thus forming the ideal negative control for optimizing the m6A-ELISA protocol (Schwartz et al. 2013a). Additionally, we used *in vitro* transcription of mRNA to generate negative and positive controls that contain either only unmodified adenosines or only m6A modified adenosines. We first assessed whether differences in m6A levels could be detected using the ABClonal-A19841 antibody (Figure 1A). Both *in vitro* transcribed m6A containing mRNA and mRNA isolated from wild-type cells showed at least 2.5-fold enrichment over controls (*in vitro* transcribed unmodified RNA or *ime4*Δ).

**Figure 1.**
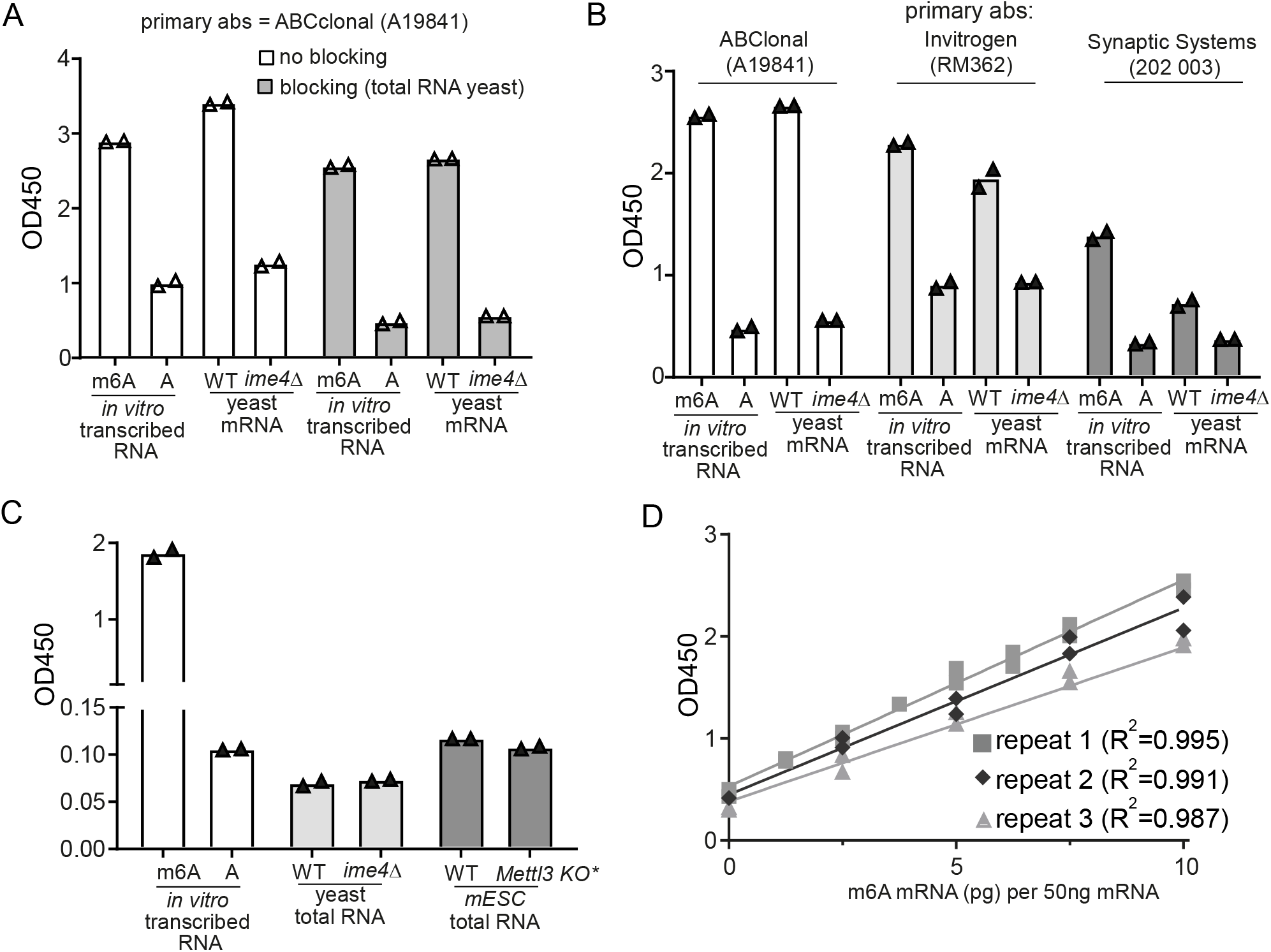
Optimization of m6A-ELISA method. **(A)** Primary antibody and blocking reagent testing. Primary antibody ((ABClonal-A19841) was incubated with or without 0.5 ug/mL of total RNA to reduce background binding to unmodified RNA. We used 50ng *in vitro* transcribed RNA generated with unmodified adenosine (A), and 0.01ng of *in vitro* transcribed m6A containing RNA (m6A). Additionally, we used polyA-selected mRNA samples from wild-type (WT) (FW1511) diploid cells and cells harbouring gene deletion in *IME4* (*ime4*Δ) (FW7030) that were induced to enter meiosis. (**B**) Primary antibody testing of samples using m6A-ELISA. Two other commercially available m6A antibodies (Invitrogen-RM362 and Synaptic systems 202003) were tested in the m6A-ELISA. (**C**) Similar analysis as in A, except that isolated total RNA was used for the analysis for WT and *ime4*Δ cells induced to enter meiosis. Also, we used total RNA isolated from mESC cells and a *Mettl3* knockout (KO). * the mESC line also contained an uninduced copy of human *Mettl3* under the control of a doxycycline inducible promoter. **(D)** Standards generated from serial dilutions and mixing of *in vitro* transcribed RNA with m6A modified adenosine and unmodified adenosine. Each standard dilution contained 50ng unmodified *in vitro* RNA with different quantities (0-10pg) of m6A modified *in vitro* transcribed RNA.

A common problem of antibody-based methods is nonspecific binding of the primary antibody to samples, which causes variability in background levels. Nonspecific binding of anti-m6A antibodies to RNA has been detected when *ime4*Δ was used as the negative control in anti-m6A mRNA pulldown followed by sequencing (Schwartz et al. 2013a). To reduce the unspecific background binding of the primary antibody, we used total RNA from *ime4*Δ cells as a blocking reagent when we incubated the primary antibody with samples (Figure 1A). The background binding to *in vitro* transcribed unmodified RNA was significantly reduced when competing total RNA was added (Supplementary Figure 1A). It should be noted that the background signal was dependent on the presence of RNA bound to wells and not due to unspecific binding of the primary antibody alone (Supplementary Figure 1B). We also tested yeast tRNA as a possible blocking reagent, but this did not result in lower background binding (Supplementary Figure 1B). We also tested two other anti-m6A antibodies (Invitrogen-RM362 and Synaptic systems 202003) under the same conditions (Figure 1B). Both antibodies showed enrichment for both the *in vitro* transcribed m6A RNA and yeast m6A mRNAs compared to unmodified RNA controls (*in vitro* transcribed unmodified RNAs, and *ime4*Δ). However, the highest signal-to-noise ratio was observed with the A19841 antibody for both *in vitro* transcribed RNA and yeast mRNA (Figure 1B). This prompted us to use the A19841 antibody for the subsequent experiments.

M6A occurs primarily on mRNAs, but it is also present on ribosomal RNA (Maden 1986; Zaccara et al. 2019; Ignatova et al. 2020; Leismann et al. 2020; Mirza et al. 2022). Hence, we typically perform m6A based assays on polyadenylated (polyA) purified mRNAs. However, some commercially available m6A assays claim that total RNA can used for the analysis (see: Materials and Methods). This prompted us to test whether total RNA isolated from yeast and mESCs is suitable for detecting m6A by m6A-ELISA. We observed no enrichment in m6A signal in WT total RNA samples compared to *ime4*Δ, likely because m6A levels were too low to be detected (Figure 1C). Total RNA m6A signal was comparable to the *in vitro* transcribed RNA with unmodified A, suggesting that there is no detectable m6A on yeast total RNA. Increasing the amount of total RNA did not improve the ability to detect m6A levels, but instead increased the variability between technical replicates, potentially due to oversaturation of the wells (Supplementary Figure 1C). Furthermore, total RNA isolated from mESCs also showed little m6A enrichment compared to the *in vitro* transcribed RNA with unmodified A, suggesting m6A levels on ribosomal RNA are below the detection range of the antibody. In line with the yeast data, we observed little difference when comparing the WT mESC to a *Mettl3* knockout. For clarity, the *Mettl3* knockout cell line also contained an uninduced copy of human *METTL3* under a doxycycline inducible promoter. We conclude that m6A-ELISA works on polyA-selected mRNAs but cannot detect m6A samples from isolated total RNA.

Next, we introduced a standard for the m6A-ELISA by making serial dilutions of *in vitro* transcribed m6A labelled RNAs. Given that the antibody has an affinity for unmodified RNA, albeit much weaker compared to m6A-containing RNA, we kept the amount of RNA for each sample constant. Hence, we generated serial dilutions using *in vitro* transcribed unmodified RNA mixed with various amounts of *in vitro* transcribed m6A modified RNA, typically 50 ng in total. We tested the standard curve linearity and reproducibility from three independent experiments. We found that the standards gave linear slopes (R^2^ = 0.99) and were reproducible (Figure 1D).

The critical steps of the m6A-ELISA protocol are summarized in Figure 2. First, total RNA is purified from cells and mRNA is subsequently isolated by two rounds of polyA purification. The purified mRNAs are quantified and incubated with binding solution to allow RNA binding to wells. We typically include positive and negative controls (WT and *ime4*Δ) and standards of *in vitro* transcribed m6A RNAs. The next steps include primary antibody incubation plus blocking RNA, washes, secondary antibody incubation, washes, and incubation with substrate solution. Stop solution is added after the positive controls have developed a medium blue colour, after approximately 10 to 30 minutes. For the analysis, we compute a standard curve from the *in vitro* transcribed m6A RNA serial dilutions, which we use to normalize the absorbance signal of samples. The m6A-ELISA assay has the potential to give absolute quantitative signal if the number of As in the standards are well defined and are spaced within the transcript for antibody accessibility. However, we did not consider this for the standard we used. In addition, while the relative differences and trends among samples were reproducible, we observed that samples and standards can give some level of variability between different ELISA experiments. Therefore, it is important to include samples that are directly compared to each other in the same m6A-ELISA experiment. Ideally, positive and negative control samples should be included, e.g. WT and *ime4*Δ. The m6A-ELISA protocol thus allows for rapid measurement of relative m6A abundance in mRNA samples.

**Figure 2.**
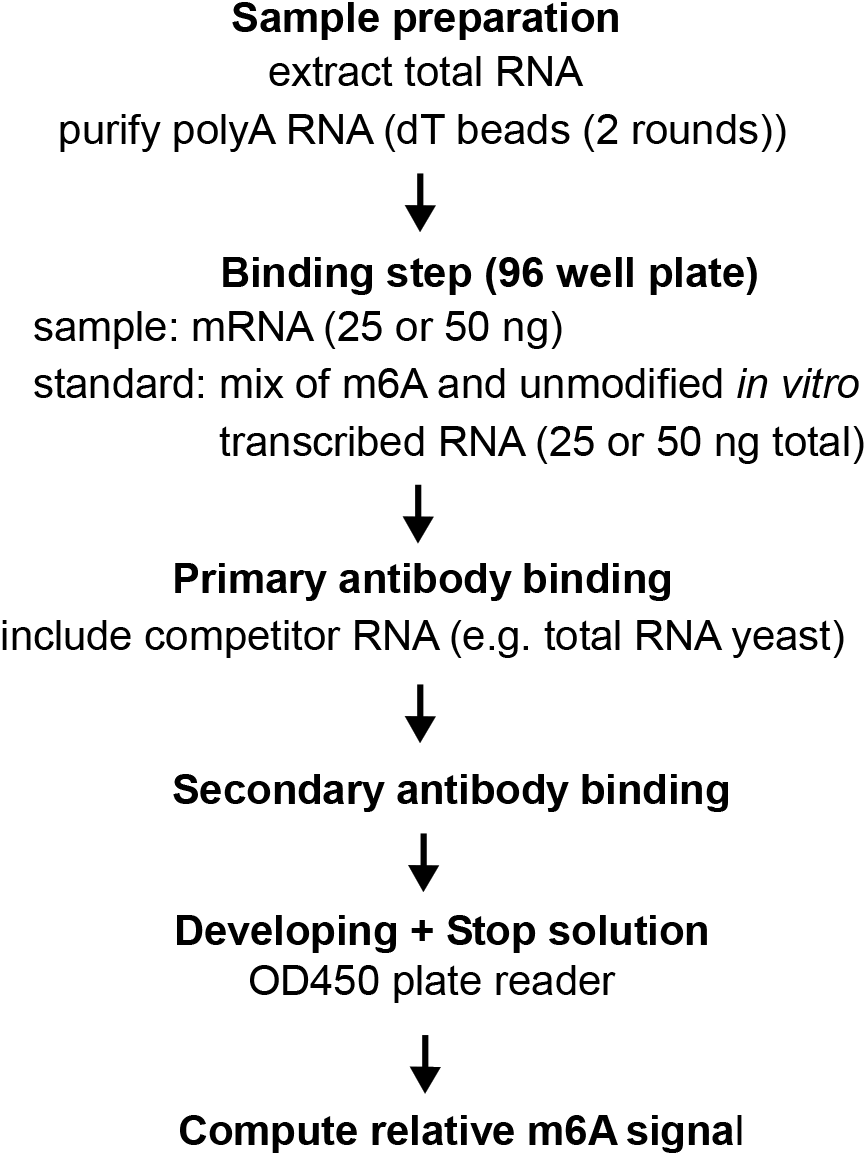
Overview of the m6A-ELISA protocol.

### Validation of the method using yeast and mammalian mRNA

Next, we validated the m6A-ELISA method by mixing different ratios of yeast WT and *ime4*Δ mRNAs. The signal increased with the relative amount of m6A-containing wild type mRNA (Figure 3A). As expected, when we used mRNAs from *ime4*Δ cells only, the normalized signal was close to background levels. Additionally, when we made 25% incremental increases in WT versus *ime4*Δ mRNA ratios, the m6A signal increased proportionately. Thus, the m6A-ELISA method can detect 25% reduction in m6A levels, as each 25% decrease was significant.

**Figure 3.**
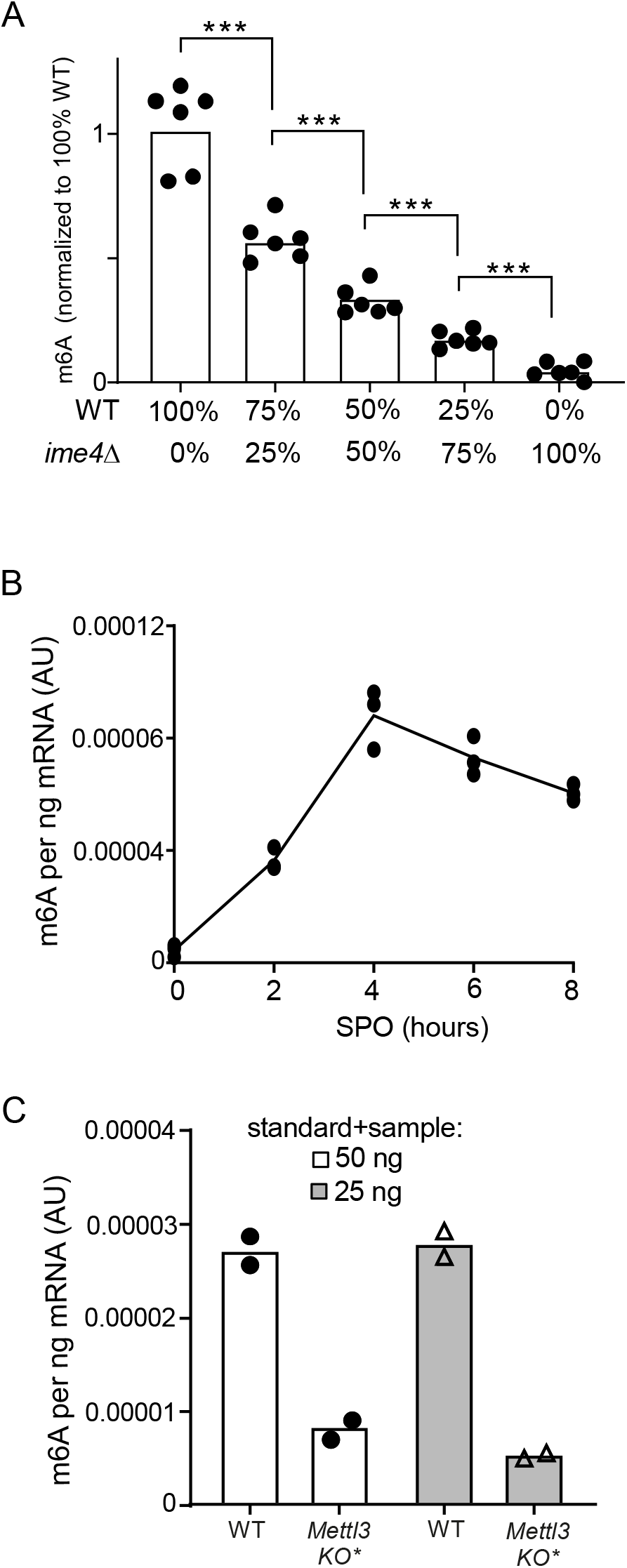
Validation of m6A-ELISA protocol. **(A)** Quantification of m6A RNA levels in WT and *ime4*Δ. WT and *ime4*Δ samples were mixed in various proportions: 100% vs 0%, 75% vs 25%, 50% vs 50%, 25% vs 75% and 0% vs 100% (WT vs *ime4*Δ). Signals were normalized to standard curve, and subsequently the WT signal was set to 100%. Unpaired *t*-test, 6 technical replicates in 2 independent experiments, each step *P* < 0.0005. **(B)** m6A deposition throughout yeast meiosis. Samples were taken at the indicated time points, and m6A levels were determined. Signals were normalized to standard curve. **(C)** m6A levels determined by m6A-ELISA using mRNA isolated from WT and *Mettl3* KO mESC. * the mESC line also contained an uninduced copy of human METTL3 under controls of a doxycycline inducible promoter. 50ng and 25ng of RNA (sample and standard) was used for the m6A-ELISA, respectively.

Commercial ELISA-based assays for m6A are available. We performed several experiments, but found that results varied strongly between lot numbers. When using purified mRNAs, some experiments showed enrichment for the WT compared to the *ime4*Δ, while in other experiments no signal over background was observed using the same samples (WT vs *ime4*Δ, Supplementary Figure 1D, representative data shown). Consistent with our data, the commercial kit also gave little signal for the WT compared to *ime4*Δ when using total RNA. This data suggests that while the commercially available m6A-ELISA assay could give a high signal-to-noise ratio on purified mRNA, it cannot be used for (yeast) total RNA, and the result can be highly variable between different lot numbers (Supplementary Figure 1D).

To validate the m6A-ELISA protocol further we determined m6A dynamics in yeast and m6A levels in mESC lines. In budding yeast, m6A rapidly increases in early stages of meiosis and subsequently decreases as cells progress further and form spores (Agarwala et al. 2012; Schwartz et al. 2013b). To test whether m6A-ELISA can detect the dynamics of m6A deposition, we took mRNA samples throughout meiosis. We found that m6A levels increased and decreased in line with what has been described previously, indicating that m6A-ELISA is suitable for detecting incremental changes in m6A levels (Figure 3B) (Agarwala et al. 2012; Schwartz et al. 2013a). Also, we measured relative m6A levels in WT mESC and a *Mettl3* knockout mESC line (Figure 3C). M6A levels were reduced over 3-fold in the *Mettl3* knockout cell line, which is consistent with the idea that METTL3 is the major m6A methyltransferase for mRNAs in mammals (Poh et al. 2021). Furthermore, we found that low technical variability could be achieved when using as little as 25 ng mammalian mRNA (Figure 3C and Supplementary Figure 2). Noteworthy, we detected leaky expression of human METTL3 from the doxycycline inducible promoter in the cell line, which may contribute the residual signal detected (Figure S3A and S3B). We conclude that m6A-ELISA can detect changes in m6A levels in purified mRNAs both in yeast and mammals.

In summary, we have developed an m6A-ELISA assay for quick and cost-effective detection of relative m6A levels from mRNA samples. For the m6A-ELISA protocol we identified and optimized critical steps that improved the signal-to-noise ratio for detection. Noteworthy, we recently applied the m6A-ELISA to measure m6A relative mRNA decay rate (Varier et al. 2022). The m6A-ELISA method could also be adapted to other mRNA modifications, provided specific antibodies are available. The m6A-ELISA is a quick, easy, and accessible detection method for global m6A detection. We propose that m6A-ELISA can be used to rapidly measure relative m6A levels across different cell types or mutants, as well as to determine the dynamic changes of m6A levels over time.

## Materials and Methods

### Yeast strains and grown conditions

Entry into meiosis was induced in the yeast SK1 diploid cells as previously described, following a standard protocol (Moretto et al. 2021). The wild-type and *ime4*Δ strains (FW1511 and FW7030) were previously described (Varier et al. 2022). In short, cells were grown at 30°C to saturation in YPD [1% (w/v) yeast extract, 2% (w/v) peptone, 2% (w/v) glucose supplemented with 24 μg/mL uracil and 12 μg/mL adenine], then diluted at OD_600_ = 0.4 to pre-sporulation medium BYTA [1.0% (w/v) yeast extract, 2.0% (w/v) bacto tryptone, 1.0% (w/v) potassium acetate, 50 mM potassium phthalate] and grown for 16-18 hours, and finally transferred at OD_600_ = 1.8 to sporulation medium SPO [0.3% (w/v) potassium acetate and 0.02% (w/v) raffinose]. Cell pellets for RNA extraction were collected 4 hours after transfer to sporulation medium.

### Embryonic stem cell culture

Mouse ESC IDG3.2 (PMID: 31047794) were maintained 2i+LIF conditions in Neurobasal (21103049)-DMEM/F-12 (11320033) 1:1 medium containing 0.5% N2 (17502048) and 1% B27 (17504044) supplements, 0.05% BSA (A7979, Sigma Aldrich), 1% Glutamax (35050), 1% nonessential amino acids (1140050) and 50μM 2-mercaptoethanol (31350-010) (all Thermo Fisher Scientific unless specified), 1,000 U/ml LIF (ESGRO ESG1107, Merck), with additional use of small molecule inhibitors: 1μM PD0325901 (04-0006, Stemgent) and 3μM CHIR99021 (1386 Axon Medchem). Cells were passaged using Stempro-Accutase (A1110501, Thermo Fisher Scientific) and grown on 0.1% gelatin (G1393, Sigma Aldrich)-coated plates.

### Generation of CRISPR/Cas9 genome engineered *Mettl3* KO mESC lines

To generate Mettl3 knockout cells, we used single gRNAs with target sites within the sixth Mettl3 exon to introduce an out-of-frame point mutation. gRNA oligos were cloned into the SpCas9-T2A-PuroR/gRNA vector (px459) via cut-ligation (gRNA sequence TTGTGATGGCTGACCCACCT). mESCs were transfected with an equimolar amount of each gRNA vector. Two days after transfection, cells were plated at clonal density and subjected to a transient puromycin selection (1 μg/mL) for 40 h. Colonies were picked 6 days after transfection and PCR primers were used to identify clones in which the sixth exon had been modified. This was confirmed with Sanger sequencing and loss of METTL3 expression was assessed by western blot (Supplementary Figure S3A). The western blot was probed for METTL3 (Abcam, ab195352), Mettl14 (Abcam, ab264408), and GAPDH (Abcam, ab8245).

### Generation of inducible FLAG-METTL3 ESC line

To generate the inducible PiggyBac donor vector with an N-terminal FLAG tag fused to METTL3, we first synthesized gBlock (IDT, Coralville, IA, USA) containing the BamHI-1XFLAG-METTL3-BamHI sequence. This fragment was then cloned into the BamHI entry site in the enhanced piggyBac transposable vector epB-Bsd-TT via cut-ligation to generate the vector, ePB-1xFLAG-METTL3 (Rosa et al. 2014).

Mettl3 KO ESCs were transfected with an equimolar amount of ePB-1xFLAG-METTL3 and piggyBac transposase (Rosa et al. 2014). Two days after transfection, cells were plated at clonal density and subjected to a transient puromycin selection (1 μg/mL) for 7 days. Colonies were picked 12 days after transfection and western blot against endogenous METTL3 (Abcam, ab195352) was performed to identify clones which maintain comparable METTL3 expression between wild type ESC cells and ePB-METT3 ESC line that was generated from parental Mettl3 KO ESCs. For experiments involving the ePB-METT3 ESC line, all inductions were performed using 100 ng/mL doxycycline (Sigma-Aldrich). A western blot of induced and uninduced cells, probed for METTL3 (Abcam, ab195352), and histone H3 (Abcam, ab1791), is shown in Supplementary Figure 3B.

### RNA extraction

Total RNA was extracted from frozen yeast pellets as previously described, using Acid Phenol:Chloroform pH 4.5 and Tris-EDTA-SDS (TES) buffer (0.01 M Tris-HCl pH 7.5, 0.01 M EDTA, 0.5% w/v SDS). The mixture was incubated at 65 °C for 45 minutes (1400RPM), transferred to ice and centrifuged at 16000g 4°C for 10 minutes. The aqueous phase was transferred to ethanol with 0.3 M sodium acetate and RNA was precipitated at −20 °C overnight. After centrifugation (16000g, 30 minutes) and washing with 80% (v/v) ethanol solution, dried RNA pellets were resuspended in RNase-free water (Ambion). Samples were further treated with rDNase (cat no 740.963, Macherey-Nagel) and column purified (cat no 740.948, Macherey-Nagel), following manufacturer’s protocols. RNA from mESCs was extracted using the RNeasy Mini Kit (cat no 74104, Qiagen) with on-column rDNase digestion (79254, Qiagen) according to manufacturer’s protocol.

### PolyA selection

Total RNA was enriched for polyA+ RNA using Oligo(dT)25 Dynabeads (Invitrogen 61005) according to manufacturer’s protocol. Two consecutive rounds of purification were performed, an initial round with 75ug total RNA and 200uL bead solution, and a second round with the eluted RNA and 40uL bead solution.

### *In vitro* transcription for generating standards

Standards were generated using the MEGAscript T7 Transcription Kit (Invitrogen AM1333), following the manufacturer’s instructions and using the included control template. Reactions either used the included ATP nucleotide solution, or N6-Methyl-ATP (Jena Bioscience NU-1101L) at the same concentration. For a 50ng standard, 50ng A containing RNA was supplemented with 0-15 pg of m6A containing RNA.

### ELISA assay

RNA concentrations were determined using the Qubit™ RNA High Sensitivity (HS) kit. 90ul of binding solution (ab156917) was added to a clear 96 well plate (ab210903) and mixed with the desired amount of mRNA sample (typically 50 ng or 25 ng). Standards containing the same quantity of *in vitro* transcribed RNA were used. The plate was incubated at 37°C for 2h, and each well was washed 4 times with PBST (0.1%), and 100uL of primary antibody solution was added and incubated for 1h at room temperature (1:10000 Abclonal A19841 including 0.5ug/mL competing *ime4*Δ yeast total RNA). Each well was then washed 4 times, incubated with secondary antibody solution for 30 minutes at room temperature (1:5000 ab205718), and washed 5 times. 100ul TMB ELISA Substrate (Fast Kinetic Rate) (ab171524) was added, and the samples were developed for up to 30 minutes. The reaction was stopped by the addition of 100uL stop solution (ab171529). The absorbance was read at 450nm using a Tecan Sunrise plate reader and Magellan software. A detailed step-wise protocol is included in Supplementary file 1. The m6A RNA methylation quantification kit from EpiGentek (P-9005) was performed according to manufacturer’s protocol, except 100ng mRNA was used instead of the recommended total RNA in designated samples.

## Author contributions

F.J.v.W. and I.E. conceived the project. I.E. performed most experiments. D.S. was involved in the initial set up of the protocol and performed experiments with commercial m6A-ELISA kits. M.M. generated the *Metll3* KO + transgene mESC line. C.C. and P.T. confirmed mESC lines by western blot. F.J.v.W. and I.E. wrote the manuscript with input from the other authors. F.J.v.W. supervised the project.

## Acknowledgements

We are grateful to Jernej Ule for suggestions, specifically with the mESC part of the manuscript. We thank the members of the van Werven lab for critical reading of the manuscript. This research was funded in whole, or in part, by the Wellcome Trust (FC001203). For the purpose of Open Access, the author has applied a CC BY public copyright licence to any Author Accepted Manuscript version arising from this submission. This work was supported by the Francis Crick Institute (FC001203), which receives its core funding from Cancer Research UK (FC001203), the UK Medical Research Council (FC001203), and the Wellcome Trust (FC001203).

---

**Supplementary Figure 1.**

**(A)** Blocking reagent testing. Primary antibody ((ABClonal-A19841) was incubated with or without 0.5 ug/mL of total RNA to reduce background binding to unmodified RNA. We used 50ng polyA-selected mRNA samples from wild-type (WT) (FW1511) diploid cells and cells harbouring gene deletion in *IME4 (ime4Δ*) (FW7030) that were induced to enter meiosis, and normalised samples to the WT signal. Background signal in *ime4*Δ RNA samples is significantly reduced after blocking (unpaired t-test of 4 technical replicates across two independent experiments, *P* = 0.0004). **(B)** Comparison of different blocking reagents. Primary antibody ((ABClonal-A19841) was incubated with no blocking RNA, with 0.5 μg/mL of total RNA or 5 μg/mL tRNAs to reduce background signal. We used polyA-selected mRNA samples from wildtype (WT) (FW1511) diploid cells and cells harbouring gene deletion in *IME4* (*ime4*Δ) (FW725) that were induced to enter meiosis. We also determined the m6A-ELISA signal of wells with no sample (-). (**C**) m6A-ELISA on samples isolated from total RNA. Total RNA was used for the analysis of samples isolated from WT and *ime4*Δ cells induced to enter meiosis. 50 ng and 500 ng of total RNA was used for the analysis. **(D)** m6A-ELISA using the EPIGENTEK kit. We performed m6A-ELISA on total RNA and mRNA purified samples using two different lot numbers. Also included are the positive (0.01ng) and negative controls from the EPIGENTEK kit.

**Supplementary Figure 2. (A)** Quantification of m6A RNA levels in WT and *ime4Δ*. Signals were normalized to standard curve. WT and *ime4*Δ samples were mixed in various proportions: 100% vs 0%, 75% vs 25%, 50% vs 50%, 25% vs 75% and 0% vs 100% (WT vs *ime4Δ*). **(B)** m6A levels determined by m6A-ELISA, but not normalized to a standard. We used mRNA isolated from WT and *Mettl3* KO mESC. * the mESC line also contained an uninduced copy of human *Mettl3* under control of a doxycycline inducible promoter. 50ng and 25ng of RNA was used, respectively.

**Supplementary Figure 3. (A)** Western blots of WT and Mettl3 KO mESC.

Membranes were probed for METTL3 (Abcam, (ab195352), Mettl14 (Abcam, ab264408), and GAPDH (Abcam, ab8245). (**B**) Western blot of WT and Mettl3 KO harbouring the human METTL3 under control of doxycycline inducible promoter mESC. Cells were grown in 2iLIF in the absence or presence of 100ng/ml doxyclyine. Membranes were probed for METTL3 (Abcam, (ab195352), and histone H3 (Abcam, ab1791). * Mettl3 KO mESC line also harbouring human METTL3 under controls of a doxycycline inducible promoter.

