## Supplementary figures and images for "m6A-ELISA, a simple method for quantifying *N6*-methyladenosine from mRNA populations"

### Supplementary figures

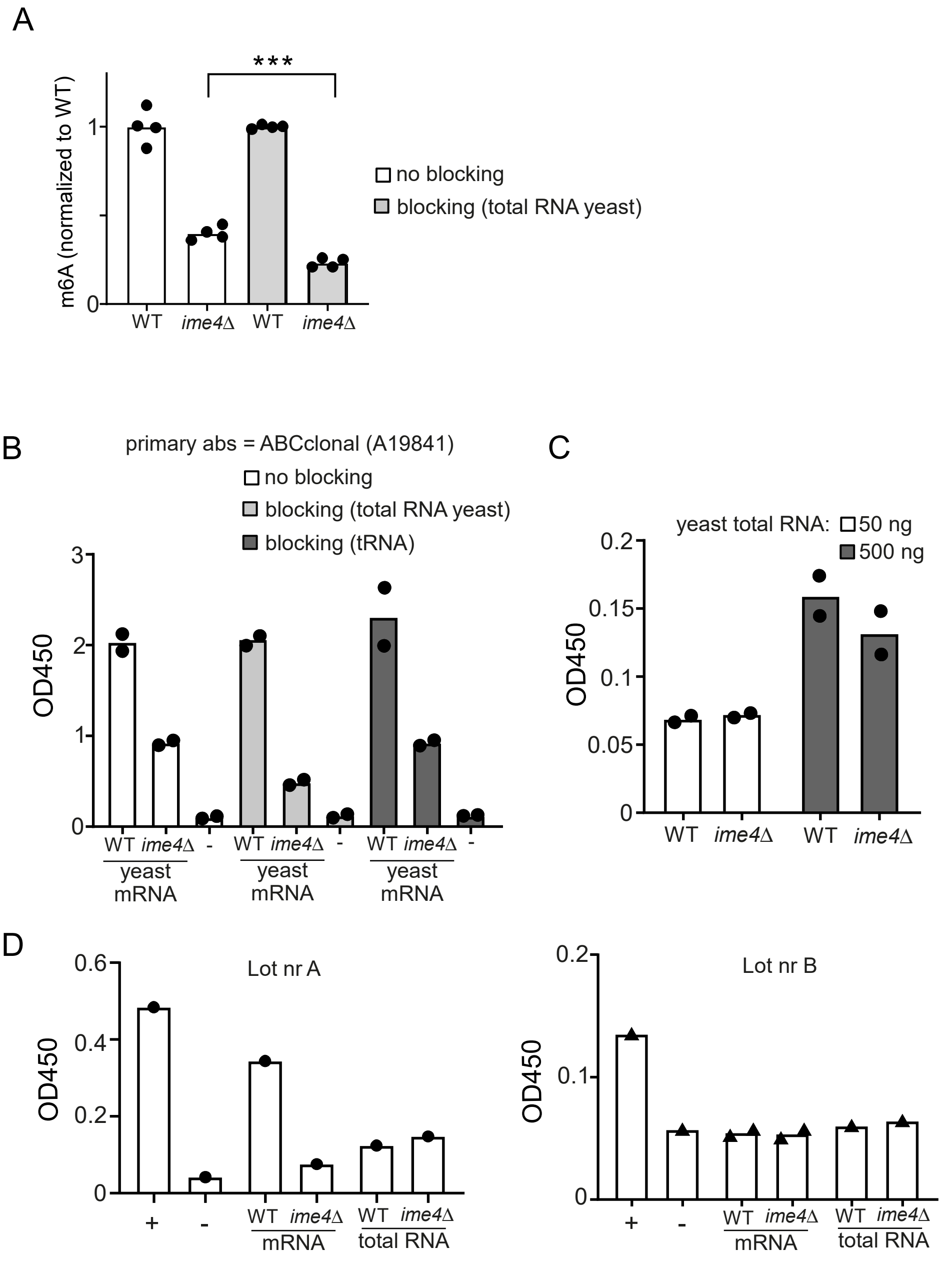

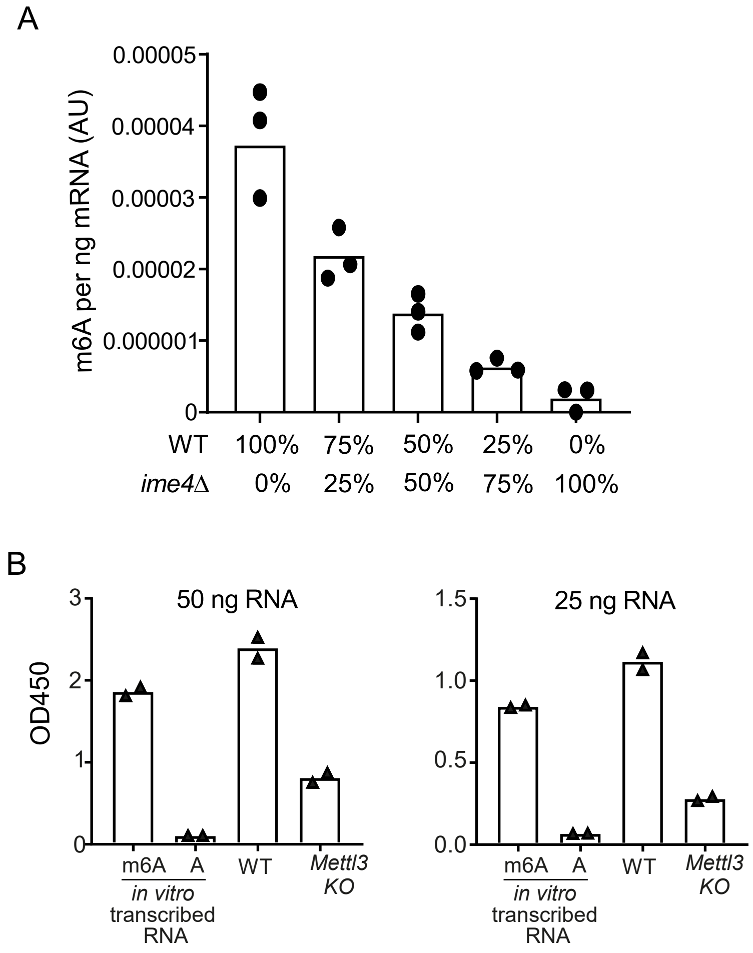

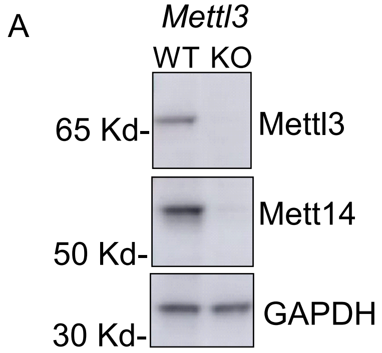

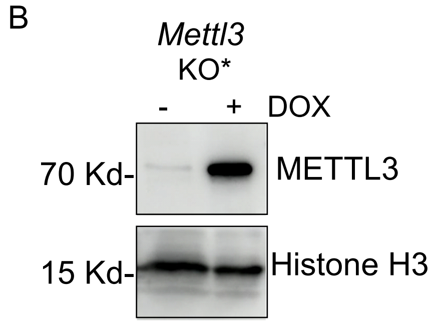
