## Supplementary file 1 for "m6A-ELISA, a simple method for quantifying *N6*-methyladenosine from mRNA populations"

**Protocol for ELISA:**

1. Add 90 ul binding solution (ab156917) to each well.

2. Add 50ng mRNA to each well. Use at least duplicates. *mRNA is obtained from total RNA by two rounds of PolyA enrichment using Oligo-dT Dynabeads. Concentrations are determined using Qubit RNA High Sensitivity kit.*

3. Incubate at 37C for 2h.

4. Prepare the primary antibody solution (mRabbit A19841, 1:10000 in PBST) with 0.5 ug/mL total RNA from *ime4Δ* (7030) strain. 100uL is needed for each well. *Adding competing yeast total RNA (without m6A) to the antibody solution improves the signal to noise ratio without causing any background signal.*

5. Remove binding solution with multichannel and wash each well with 4x 200uL PBST (0.1%).

6. Add 100ul of primary Ab dilution (*step 4*) to each well. Make sure to remove any bubbles. Incubate for 1h at room temperature.

7. Remove and wash 4x with 200uL PBST.

8. Add 100uL secondary antibody solution (anti-rabbit ab205718 1:5000 in PBST). Make sure to remove any bubbles. Incubate for 30 minutes at room temperature.

9. Remove and wash 5x with 200uL PBST. After final wash, make sure to tap plate onto tissue to remove as much of the PBST as possible.

10. Add 100 ul developing ('fast kinetic rate' ab171524) solution. Make sure to remove any bubbles. Keep covered from light. *Optional: The development can be measured on plate reader at 650+450nm.*

11. When satisfied with development, stop the plate reader and add 100uL stop solution. *Differences in signal can often be observed within a few minutes, but I found that a longer development of up to 30 minutes works well for my samples.*

12. Read absorbance at 450 nm.

**Appendix: Standard curves for ELISA.**

The standard curve contains *in vitro* transcribed mRNA without m6A (‘A’) mixed with small amounts of *in vitro* transcribed mRNA using only m6ATP and no ATP (‘m6A’). The A only control, using an equal amount of mRNA as the samples, is crucially important as it allows to correct for background binding of the antibody to RNA.

*Instead of a full standard curve, a positive (A+m6A) and negative (A only) control could be used.*

Suggested amounts:

50ng A + 0.015ng m6A

50ng A + 0.01 ng m6A

50ng A + 0.0075 ng m6A

50ng A + 0.005 ng m6A

50ng A + 0.00375 ng m6A

50ng A + 0.0025 ng m6A

50ng A + 0.00125 ng m6A

50ng A

To make the standard curve

- Make dilutions of m6A *in vitro RNA*
- Dilute 1:10 m6A RNA down to 1ng/uL
- Dilute 1ng/uL to 0.02ng/uL (98uL H2O + 2uL stock)
- Dilute further:

| standard | uL 0.02ng/uL stock | uL H2O |
| --- | --- | --- |
| 0.015ng/uL | 15 | 5 |
| 0.01ng/uL | 10 | 10 |
| 0.0075ng/uL | 7.5 | 12.5 |
| 0.005ng/uL | 5 | 15 |
| 0.00375ng/uL | 3.75 | 16.25 |
| 0.0025ng/uL | 2.5 | 17.5 |
| 0.00125ng/uL | 1.25 | 18.75 |

- Measure concentration of A *in vitro* RNA and determine amount for 50ng
- Add 90ul binding solution to each well, then add 50ng in vitro A RNA to each well, then add 1uL of each standard

**Materials and reagents**

Materials ELISA:

- 96-Well Microplate Pack (ab210903)
- DNA Binding Microplate Solution (ab156917)
- N6-methyladenosine / m6A Rabbit mAb (A19841)
- Phosphate buffered saline (PBS) + 0.1% TWEEN-20 (9416) (PBST)
- Goat Anti-Rabbit IgG H&L (HRP) (ab205718)
- TMB ELISA Substrate (Fast Kinetic Rate) (ab171524)
- 450 nm Stop Solution for TMB Substrate (ab171529)
- TECAN Sunrise Plate Reader

Generating standards:

- MEGAscript T7 Transcription Kit-25 reactions (AM1333)
- N6-Methyl-ATP (NU-1101L)

Preparing mRNA samples from total RNA:

- Qubit™ RNA High Sensitivity (Q32855)
- Dynabeads™ Oligo(dT)25 (61005)
